# Fc effector activity and neutralization against SARS-CoV-2 BA.4 is compromised in convalescent sera, regardless of the infecting variant

**DOI:** 10.1101/2022.07.14.500042

**Authors:** Simone I. Richardson, Prudence Kgagudi, Nelia P. Manamela, Haajira Kaldine, Elizabeth M. Venter, Thanusha Pillay, Bronwen E. Lambson, Mieke A. van der Mescht, Tandile Hermanus, Zelda de Beer, Talita Roma de Villiers, Annie Bodenstein, Gretha van den Berg, Marizane du Pisanie, Wendy A. Burgers, Ntobeko A.B. Ntusi, Veronica Ueckermann, Theresa M. Rossouw, Michael T. Boswell, Penny L. Moore

**Affiliations:** National Institute for Communicable Diseases of the National Health Laboratory Services, Johannesburg, South Africa; MRC Antibody Immunity Research Unit, School of Pathology, University of the Witwatersrand, Johannesburg, South Africa; Department of Immunology, Faculty of Health Sciences, University of Pretoria, Pretoria, South Africa; Tshwane District Hospital, Pretoria, South Africa; Division for Infectious Diseases, Department of Internal Medicine, Steve Biko Academic Hospital and University of Pretoria, Pretoria, South Africa; Institute of Infectious Disease and Molecular Medicine, University of Cape Town, Cape Town, South Africa; Division of Medical Virology, Department of Pathology; University of Cape Town, Cape Town, South Africa; Wellcome Centre for Infectious Disease Research in Africa, University of Cape Town, Cape Town, South Africa; Department of Medicine, University of Cape Town and Groote Schuur Hospital, South Africa; Hatter Institute for Cardiovascular Research in Africa, Faculty of Health Sciences, University of Cape Town, Cape Town, South Africa; Centre for the AIDS Programme of Research in South Africa, Durban, South Africa

**Keywords:** Omicron BA.4, antibody dependent cellular cytotoxicity, neutralization, SARS-CoV-2, breakthrough infection

## Abstract

The SARS-CoV-2 Omicron BA.1 variant, which exhibits high level neutralization resistance, has since evolved into several sub-lineages including BA.4 and BA.5, which have dominated the fifth wave of infection in South Africa. Here we assessed the sensitivity of BA.4 to neutralization and antibody dependent cellular cytotoxicity (ADCC) in convalescent donors infected with four previous variants of SARS-CoV-2, as well as in post-vaccination breakthrough infections (BTIs) caused by Delta or BA.1. We confirm that BA.4 shows high level resistance to neutralization, regardless of the infecting variant. However, breakthrough infections, which trigger potent neutralization, retained activity against BA.4, albeit at reduced titers. Fold reduction of neutralization in BTIs was lower than that seen in unvaccinated convalescent donors, suggesting maturation of neutralizing responses to become more resilient against VOCs in hybrid immunity. BA.4 sensitivity to ADCC was reduced but remained detectable in both convalescent donors and in BTIs. Overall, the high neutralization resistance of BA.4, even to antibodies from BA.1 infections, provides an immunological mechanism for the rapid spread of BA.4 immediately after a BA.1-dominated wave. Furthermore, although ADCC activity against BA.4 was reduced, residual activity may nonetheless contribute to the protection from disease.

## Introduction

The emergence of SARS-CoV-2 variants of concern (VOCs) bearing mutations in the spike protein has resulted in escape from neutralizing antibodies (mAbs) triggered by vaccination and infection (1–4), and subsequently reduced protection from infection (5). Most recently, these VOCs include Omicron BA.1, containing over 30 mutations in the spike region, against which neutralization titers are further reduced (6). In contrast, the ability of vaccines to prevent severe disease has been maintained (5,7,8). This is likely due to the preserved activity of T cells and Fc effector function, including antibody dependent cellular cytotoxicity (ADCC), against VOCs (9,10).

Omicron has since evolved into several sub-lineages including BA.2, BA.2.12.1, BA.4 and BA.5 (11). The BA.4 and BA.5 sub-lineages, which share the same spike sequence but differ from one another in non-structural protein and membrane (M) genes, drove the fifth wave of infection in South Africa, and have subsequently been detected in more than 30 other countries (12). BA.4 and BA.5 are genetically similar to BA.2 but contain two additional mutations in the receptor binding domain (RBD), L452R and F486V. As a consequence, compared to BA.1 and BA.2, BA.4 has shown increased neutralization resistance to convalescent sera, vaccinee sera and monoclonal antibodies (4,13,14).

We and others have shown that each SARS-CoV-2 variant triggers different profiles of neutralizing antibodies (nAbs) and Fc effector function (15,16). For example, the Beta variant triggered humoral responses with increased cross-reactivity, whereas Omicron triggered more strain-specific nAbs (15,16). Here, we assessed the sensitivity of BA.4 to neutralizing antibodies and ADCC elicited by infections caused by D614G, Beta, Delta or BA.1 (responsible for the first four waves in South Africa) in vaccinated and unvaccinated individuals.

We confirm that BA.4 shows high level resistance to neutralization, regardless of the infecting variant. However, high neutralizing titers associated with breakthrough infection with either Delta or BA.2 after vaccination, results in preserved neutralization against BA.4. Further, we show that while ADCC activity against BA.4 was reduced further than previously reported for other VOCs, it remained detectable in both convalescent plasma and in vaccine breakthrough infections. Overall, this study confirms the increased neutralization resistance of BA.4 and provides an immunological mechanism for the rapid spread of BA.4 in South Africa, despite high levels of infections by previous VOCs (17). Furthermore, despite the reduced ADCC against BA.4, the residual activity we detect in convalescent plasma and vaccinees may nonetheless contribute to the protection from severe disease.

## Results

### BA.4 escapes convalescent plasma neutralization, regardless of the infecting strain

We assayed plasma from individuals infected in the first four waves of infection in South Africa, with D614G (wave 1, n=16), Beta (wave 2, n=10), Delta (wave 3, n=7) or Omicron BA.1 (wave 4, n=20) with clinical and demographic details presented in **Table S1**. All samples were obtained from individuals who reported no prior infection or vaccination, confirmed by national databases (15,16). Overall, we show that BA.4 is highly resistant to neutralization, regardless of the infecting strain, with titers ranging from a GMT of 39 in D614G infections to 179 in BA.1 infection **(Figure 1)**. However, the fold loss of neutralization activity varies by wave, with Delta and BA.1 infections (both of which trigger high titers of >1:2,500 against their matched spikes, perhaps a consequence of higher viral loads) showing 34 and 17-fold losses against BA.4 respectively **(Figure 1 C, D)**. In contrast, in D614G and Beta infections, where autologous titers against the infecting strains were lower, around 1:300, the loss in neutralization against BA.4 was 5-8 fold **(Figure 1 A, B)**.

**Figure 1:**
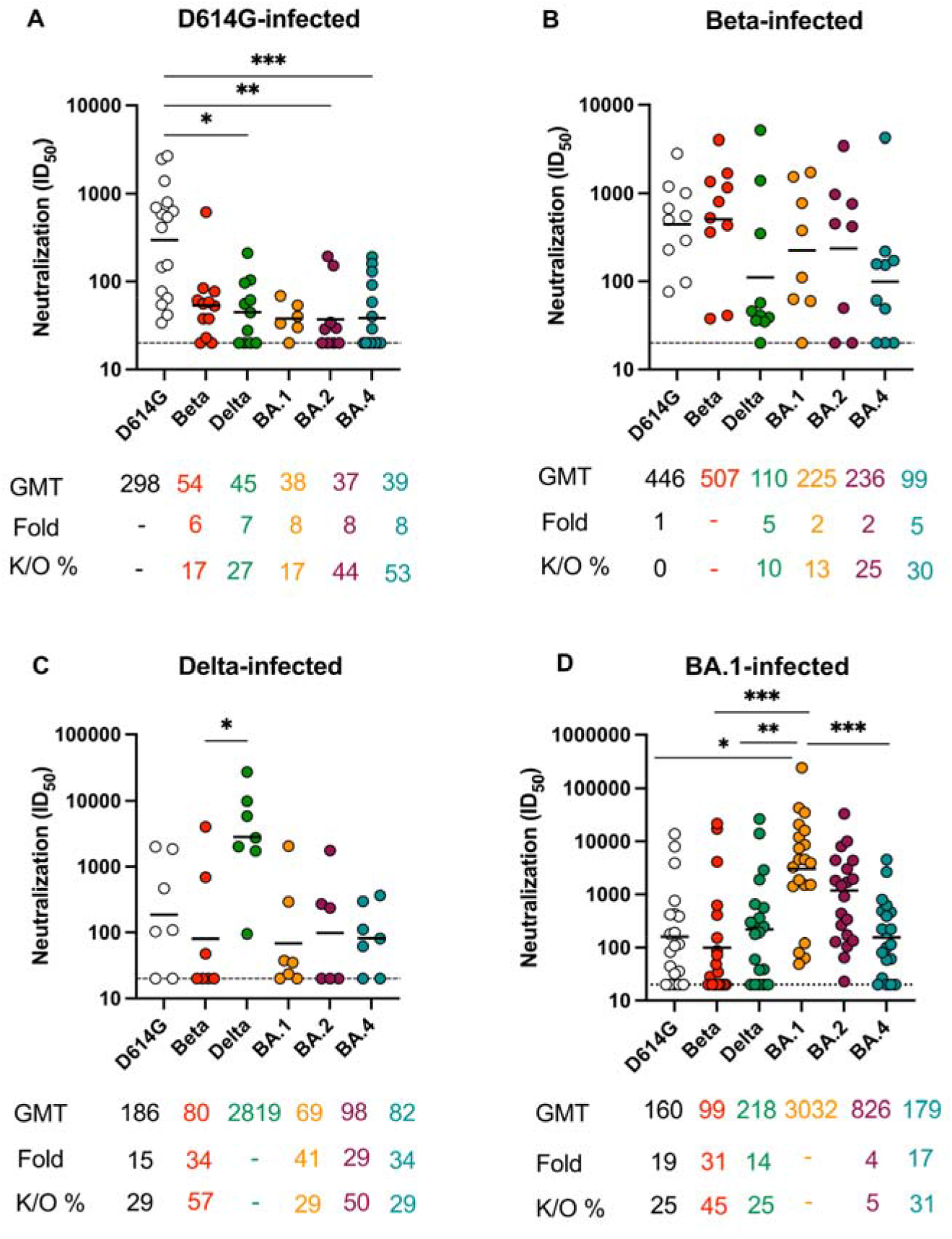
BA.4 neutralization escape varies by the infecting variant in unvaccinated convalescent individuals. Neutralization titer (ID_50_) in convalescent plasma from unvaccinated donors infected with (A) D614G, (B) Beta, (C) Delta and (D) Omicron BA.1. Plasma was tested against D614G, Beta, Delta, Omicron BA.1, BA.2 and BA.4. Lines indicate geometric mean titer (GMT) also represented below the plot with fold decrease and knock-out (K/O) of activity for other variants as a percentage relative to the infecting strain. Dotted lines indicate the limit of detection of the assay. Statistical significance across variants is shown by Friedman test with Dunn’s correction. *p<0.05; **p<0.01; ***p<0.001; ****p<0.0001 and ns = non-significant. All data are representative of two independent experiments.

We also observed variant-specific differences in neutralization of Omicron BA.2, which showed similar titers to BA.4 in D614G and Delta infection, but different titers in Beta and BA.1 infections (with BA.2 significantly more sensitive than BA.4, with a titer of 826 and 179, respectively). In general, when considering the degree of cross-reactivity of antibodies triggered by each variant against multiple VOCs, we observed a greater number of significant fold losses for antibodies triggered by D614G (significant losses against Delta, BA.2 and BA.4) and by BA.1 (with significant fold losses against all variants except BA.2), as previously reported **(Figure 1 A, D)** (16). In contrast, Beta-elicited nAbs showed greater levels of cross-reactivity than those triggered by other variants, as we have described elsewhere **(Figure 1 B)** (15,18).

### Breakthrough infection following vaccination shows increased neutralization crossreactivity against BA.4

We next tested the capacity of plasma from breakthrough infections (BTIs) caused by Delta or Omicron BA.1, following vaccination (16,19) to neutralize BA.4. We and others have previously shown that BTIs trigger high levels of nAbs that are cross-reactive for VOCs (16,19,20). In both Delta and BA.1 BTIs, titers were highest against D614G (which matches the vaccine strain), rather than the infecting variant **(Figure 2A, B**). In Delta BTIs, titers against D614G were significantly higher than against any other VOC **(Figure 2A**), whereas for BA.1 BTIs, significant fold losses compared to D614G were only observed in BA.4 **(Figure 2B**). Unlike previous VOCs, BA.4 shows substantially increased resistance to neutralization in BTIs caused by either Delta or Omicron BA.1. In Delta and BA.1 BTIs, we saw a 7-fold reduction in titers compared to titers against the infecting variant **(Figure 2A and 2B**) and in contrast to unvaccinated individuals, all samples retained neutralization activity against BA.4 **(Figure 2B**).

**Figure 2:**
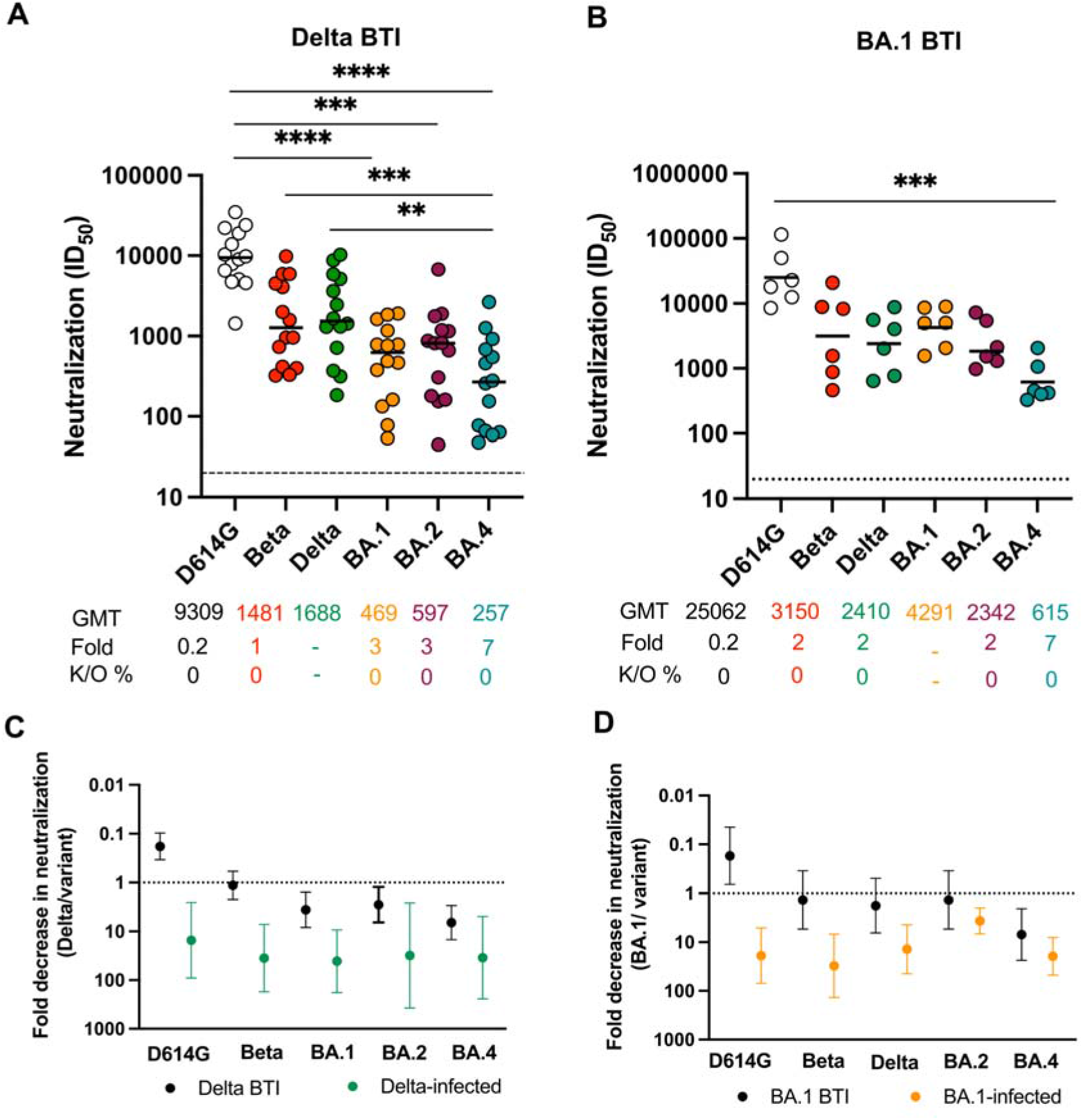
Breakthrough infections show reduced neutralization activity against BA.4, despite high titers against other VOCs. Neutralization titer (ID_50_) in convalescent plasma from vaccinated donors subsequently infected with (A) Delta and (B) Omicron BA.1. Plasma were tested against D614G, Beta, Delta, Omicron BA.1, BA.2 and BA.4. Lines indicate geometric mean titer (GMT) also represented below the plot with fold decrease and knock-out (K/O) of activity for other variants as a percentage relative to the infecting variant. Dotted lines indicate the limit of detection of the assay. Fold decrease in neutralization for each VOC represented as a ratio of the titer to the infecting variant Delta (C) or BA.1 (D), for infections in unvaccinated individuals (green for Delta and orange for BA.1) and BTIs (black). Statistical significance across variants is shown by Friedman test with Dunn’s correction. *p<0.05; **p<0.01; ***p<0.001; ****p<0.0001 and ns = non-significant. All data are representative of two independent experiments.

We next compared convalescent plasma from unvaccinated individuals to BTIs by the same variant, to assess whether similar fold losses were observed in both cases **(Figure 2C, D**). In both Delta and BA.1, enhanced titers were observed against D614G, consistent with prior exposure to the vaccine sequence (Wuhan-1), whereas all other ratios were >1, indicating decreased neutralization relative to the infecting variant. However, for Delta and BA.1, the fold decrease in neutralization against each variant was higher in unvaccinated individuals **(Figure 2C, D - green and orange**) compared to BTIs **(Figure 2C, D – black**). This suggests that two antigenic exposures result in more resilient neutralizing responses to VOCs.

### BA.4 shows increased escape from antibody dependent cellular cytotoxicity compared to other VOCs

We next assessed the ability of plasma antibodies from convalescent donors from each of the four waves to cross-link D614G, Beta, Delta, BA.1, BA.2 or BA.4 cell-surface expressed spike and activate FcγRIIIa (CD16) as a proxy for ADCC activity. As we have previously reported, fold loss in activity for ADCC was generally in the order of 2-3 fold, much less than for neutralization, likely due to the higher number of epitopes recognised (21). However, compared to ADCC against the matched spike in each wave, we observed 2- to 8.8-fold reduced activity against BA.4 **(Figure 3A-D)**. These losses were statistically significant, with the exception of Beta-triggered ADCC **(Figure 3B)**, consistent with our previous studies suggesting that Beta triggers more cross-reactive ADCC (21). Compared to BA.1, BA.4 was more resistant to ADCC in plasma from all four waves. However, despite losses, ADCC activity was retained, ranging from 628 relative light units (RLU) for D614G infections **(Figure 3A)** to 216 RLU for Delta infections **(Figure 3C)**. Thus, ADCC activity against BA.4 in convalescent plasma from unvaccinated individuals was reduced but detectable, regardless of the infecting variant.

**Figure 3:**
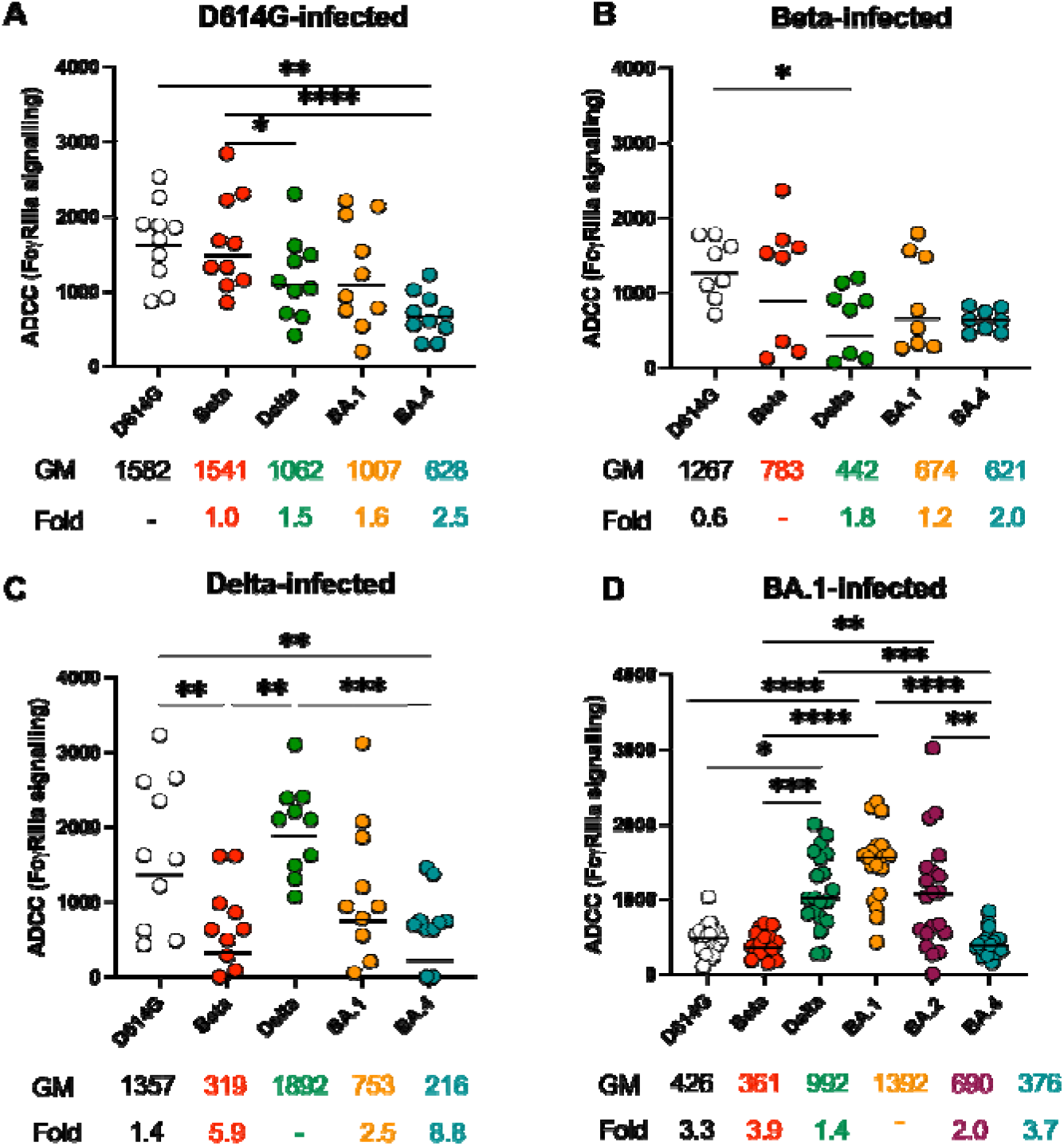
ADCC against BA.4 is reduced, but preserved in convalescent plasma from previously unvaccinated individuals, regardless of the infecting variant. Antibody-dependent cellular cytotoxicity (ADCC) in unvaccinated individuals infected with (A) D614G, (B) Beta, (C) Delta and (D) Omicron BA.1. The ability of plasma to cross-link spike expressed on the surface of HEK293T cells and activate FcγRIIIa represented as relative light units (RLU) with background as determined in the absence of antibody are shown. All data are representative of two independent experiments. Lines indicate geometric mean (GM) RLU, also represented below the plot with fold decrease of activity for other variants relative to the infecting variant. Statistical significance across variants is shown by Friedman test with Dunn’s correction and between vaccinated and unvaccinated samples by the Mann Whitney test. *p<0.05; **p<0.01; ***p<0.001; ****p<0.0001 and ns = non-significant.

### ADCC elicited by Delta and Omicron BA.1 breakthrough infections are compromised by BA.4

Using a subset of samples tested against neutralization, we measured FcγRIIIa activation for BTIs caused by Delta (n = 5) and BA.1 (n = 7) (**Figure 4A, B**). In line with what we have previously reported (16,19), ADCC activity was higher in individuals who were previously vaccinated, then infected, compared to those who were not, regardless of the infecting variant. This included higher activity against BA.4 where RLUs were 3.2 fold higher in Delta BTIs compared to Delta-infected unvaccinated individuals (**Figure 3C and 4A)** and 1.5 fold greater in BA.1 BTIs compared to BA.1-infected unvaccinated individuals **(Figure 3D and 4B)**. In contrast to neutralization, fold decreases of ADCC against VOCs relative to the infecting variant were similar in unvaccinated compared to vaccinated individuals **(Figure 4C, D)**. Regardless of the infecting variant, BA.4 showed the biggest fold decrease of ADCC in both BTIs and unvaccinated but infected individuals. This indicates that while vaccination increases ADCC activity against BA.4, it does not improve the relative cross-reactivity against this variant.

**Figure 4:**
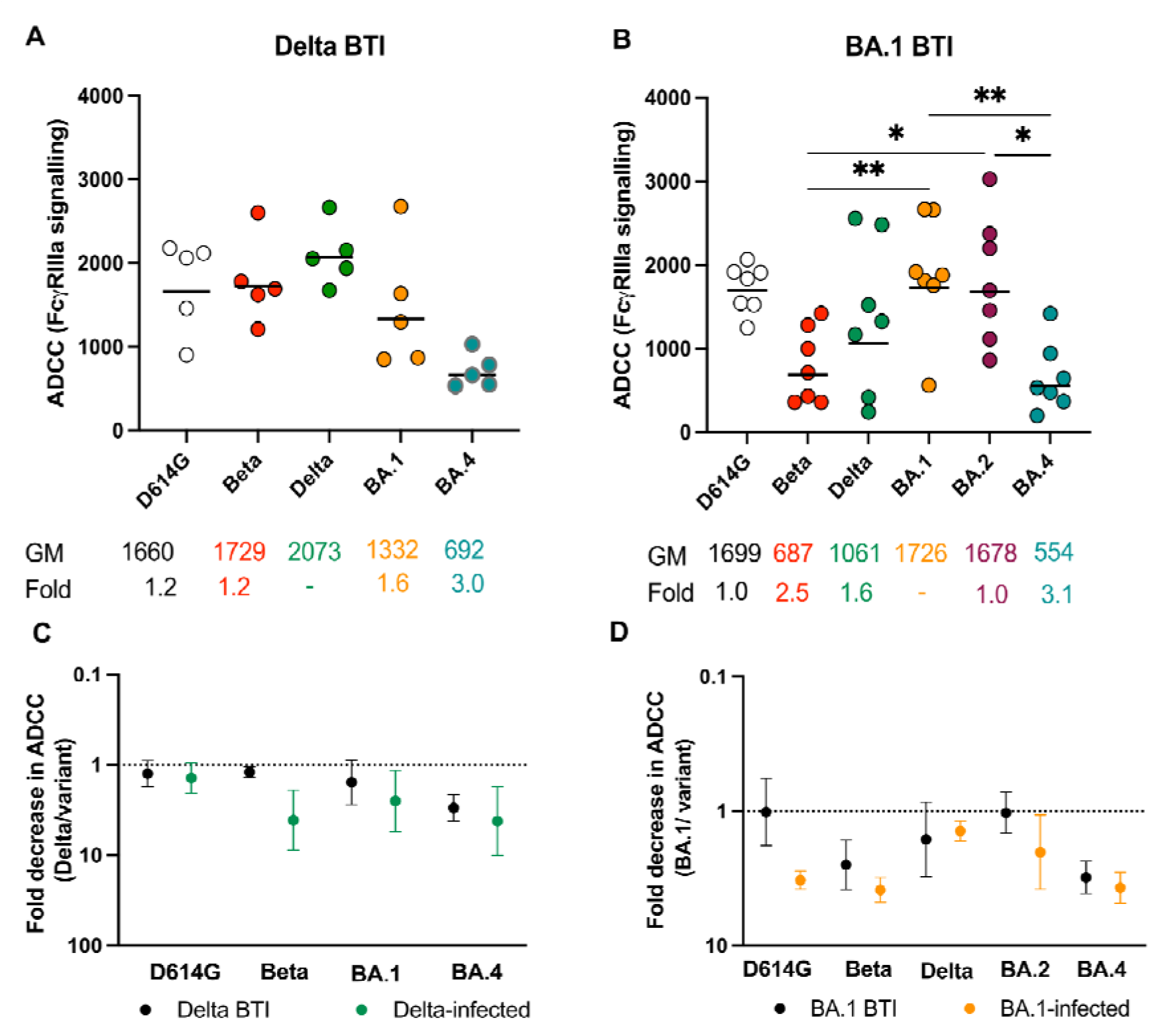
ADCC against BA.4 is reduced in BTIs caused by Delta or BA.1 to a similar extent as in convalescent plasma. Antibody-dependent cellular cytotoxicity of vaccinated donors subsequently infected with (A) Delta and (B) Omicron BA.1. The ability of plasma to cross-link spike expressed on the surface of HEK293T cells and activate FcγRIIIa represented as relative light units (RLU) with background as determined in the absence of antibody are shown. Lines indicate geometric mean (GM) RLU, also represented below the plot with fold decrease of activity for other variants relative to the infecting variant. Fold decrease in neutralization for each VOC represented as a ratio of ADCC activity to the infecting variant Delta (C) or BA.1 (D), for infections in unvaccinated individuals (green for Delta and orange for BA.1) and BTIs (black). Statistical significance across variants is shown by Friedman test with Dunn’s correction. *p<0.05; **p<0.01; ***p<0.001; ****p<0.0001 and ns = nonsignificant. All data are representative of two independent experiments.

## Discussion

The ability of BA.4 to escape neutralization elicited by vaccination and previous infection is now well described (4,14,22–25). Here, we have extended these studies to define BA.4 resistance to neutralizing and ADCC-mediating antibodies triggered by each of the four VOCs (D614G, Beta, Delta and BA.1) that sequentially caused waves of infection in South Africa (26). Regardless of the infecting variant, we show that BA.4 is highly resistant to neutralization, with particularly large reductions in neutralization for antibodies triggered by Delta and BA.1, compared to D614G and Beta. Secondly, we provide the first assessment of the effect of BA.4 mutations on Fc effector function, which has been preserved against other VOCs (21). Using FcγRIIIa activation as a proxy for ADCC, we show that BA.4 shows greater ADCC escape than previous VOCs. As for neutralization, this loss is especially pronounced in Delta and BA.1 infected individuals, including in breakthrough infections. Our data extend previous studies to assess antibodies triggered by four VOCs and confirm that BA.4 is resistant to both neutralization and ADCC, regardless of the infecting variant.

Our data confirm many studies showing that VOCs trigger responses with different specificities (15,16). Here, the largest fold decreases for neutralization against BA.4 were seen for unvaccinated individuals previously infected with Delta and BA.1 (23). For Delta, this is in contrast to a previous report, where Delta-wave patient sera neutralized not only Delta but also the BA.4/5 and BA.2.12.1 variants, which, like Delta, contain substitutions at position L452 (25). In our cohort, we noted that autologous titers to the Delta spike were higher than those in D614G and Beta, perhaps a consequence of the high viral loads that are associated with Delta infections (27). Of note, in BA.1 infections, we observed that neutralization of both BA.2 and BA.4 were reduced. However, BA.4 was significantly more resistant than BA.2, despite the fact that these two sub-lineages of Omicron are genetically very similar (11). This suggests that BA.1 triggered antibodies may target epitopes including L452 and F486 that distinguish BA.2 and BA.4, which will form the basis of future mapping studies.

Although BA.4 is highly neutralization resistant, we nonetheless observed relatively high titers in previously vaccinated individuals with BTIs. We and others have previously shown that BTIs result in significantly higher neutralization titers than in people who were not vaccinated prior to infection (16,19,23). High titers generally result in better neutralization of VOCs, which is the basis of ongoing booster regimens. However, we note that the fold loss in neutralization is higher in unvaccinated individuals compared to BTIs. This suggests that the preserved activity against VOCs such as BA.4 is not simply a consequence of higher starting titers, but that the quality of neutralizing antibodies resulting from BTI is intrinsically better. This is consistent with ongoing affinity maturation of responses after second antigenic exposures (28). In South Africa, where >95% of people are now estimated to be seropositive, this scenario of hybrid immunity is likely very common (17,29). However, even in the context of these high titer responses, BA.4 shows reduced sensitivity to neutralization compared to other VOCs, perhaps accounting for ongoing community transmission.

This study provides the first assessment of BA.4 mutations on Fc effector function (21). Here we show that BA.4 shows greater ADCC escape than previous VOCs. As for neutralization, this loss is especially pronounced in Delta and BA.1 infected individuals, including in breakthrough infections. This suggests that as for neutralization, the sequence of the infecting spike also affects the quality of antibodies mediating Fc effector function (21).

ADCC, and other Fc effector functions have proven to be remarkably resilient in the face of mutations in spike. However, our observation that BA.4 shows significantly reduced sensitivity to ADCC responses suggests limits to that tolerance and provides interesting insights into the targets of these antibodies. The observation that Beta-directed ADCC is most compromised following infection but not BTI caused by Delta suggests differences in primary versus hybrid immunity, as well as in antibodies triggered by different VOCs. Furthermore, the reduced ADCC sensitivity of BA.4 suggests that regions mutated in this VOC may define key ADCC epitopes, which may or may not overlap with sites targeted in neutralization. Delineation of these sites will be key to defining the targets of antibodies mediating ADCC. These studies will be important in the assessment of Fc effector function against emerging VOCs and inform the development of universal vaccines for improved cross-reactivity against emerging VOCs.

Overall, these data extend previous studies to assess antibodies triggered by four VOCs, and confirm that BA.4 is resistant to both neutralization and ADCC, regardless of the infecting variant. The high level of resistance of BA.4, particularly to antibodies from BA.1 infections, provides an immunological mechanism for the rapid spread of BA.4 in South Africa immediately after a BA.1-dominated wave, and provides insights into populational-level immunity gaps that may exist elsewhere. Furthermore, the reduced sensitivity of BA.4 to ADCC, unlike previous VOCs, provides useful insights for future mapping of the targets of antibodies mediating ADCC. Lastly we note that although ADCC activity against BA.4 was reduced, residual activity may nonetheless contribute to the protection from severe disease. While T cells almost certainly also contribute to this effect, the preserved ADCC against BA.4 is consistent with the observation of low levels of severe disease and hospitalisation during this wave in South Africa (30).

### Limitations of the study

We acknowledge that the numbers of individuals in several of these groups are small, and future studies should include additional donors. Additionally, not all samples were run across both ADCC and neutralization assays (as indicated in Table S1) as a result of sample availability. Furthermore, although we have extensive clinical follow-up, we cannot rule out the possibility that convalescent donors had experienced previous undocumented asymptomatic infection which could alter the quality of humoral responses. We have not included measurements of T cell responses to BA.4, which likely contribute to protection from severe disease. Lastly, viral sequences were available only for a subset of samples in each wave, though the samples were collected when each variant dominated infections during that particular wave.

## Acknowledgements

We acknowledge the participants who volunteered for this study and thank W van Hougenhouck-Tulleken for database support. We thank Rajiev Ramlall for assistance with participant recruitment at Tshwane District Hospital and Daniel Amoako and Jennifer Giandhari for sequencing support. The parental soluble spike was provided by J McLellan (University of Texas) and parental pseudovirus plasmids by Drs E Landais and D Sok (IAVI). PLM is supported by the South African Research Chairs Initiative of the Department of Science and Innovation and National Research Foundation of South Africa, the SA Medical Research Council SHIP program, the Centre for the AIDS Programme of Research in South Africa (CAPRISA). We acknowledge funding from the Bill and Melinda Gates Foundation, through the Global Immunology and Immune Sequencing for Epidemic Response (GIISER) program.

## Author contributions

S.I.R. designed the study, performed experiments, analyzed the data and wrote the manuscript. N.P.M., E.M.V. and T.P. performed Fc experiments and analyzed data. P.K., H.K. and T.H. performed neutralization assays. B.E.L. produced variant plasmids. M.A.vdM. processed samples which were recruited by V.U., T.R. and M.T.B. who established the Pretoria COVID-19 study. W.A.B. and N.A.B.N. established the Groote Schuur Hospitalised cohort. P.L.M. conceptualized the study and wrote the manuscript.

---

**Table S1:**
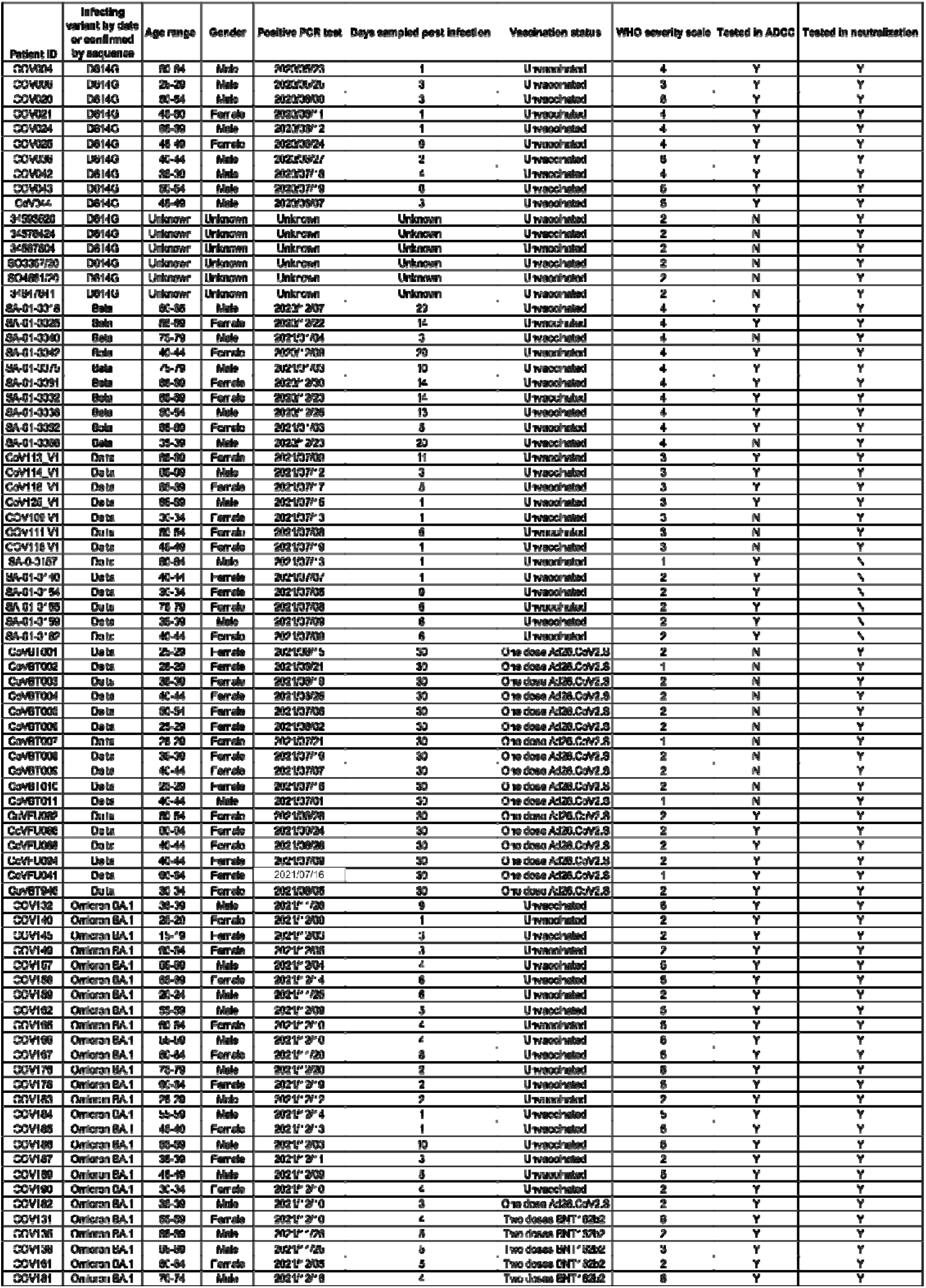
Demographic and clinical information

## Methods

### RESOURCE AVAILABILITY

#### Lead Contact

Further information and reasonable requests for resources and reagents should be directed to and will be fulfilled by the lead contact, Penny Moore.

#### Materials availability

Materials will be made by request to Penny Moore.

#### Data and code availability

■ All data reported in this paper will be shared by the lead contact upon request.
■ This paper does not report original code.
■ Any additional information required to reanalyze the data reported in this paper is available from the Lead Contact upon request.

### EXPERIMENTAL MODEL AND SUBJECT DETAILS

#### Human Subjects

Plasma samples from the first SARS-CoV-2 wave (D614G-infected) were obtained from a previously described cohort across various sites in South Africa prior to September 2020 (3). Second wave samples (Beta-infected) were obtained from a cohort of patients admitted to Groote Schuur Hospital, Cape Town in December 2020 - January 2021 (18). Third wave samples (Delta-infected) were obtained from the Steve Biko Academic Hospital, Tshwane from patients admitted in July 2021(15). Samples infected in the fourth COVID-19 wave of infection in South Africa were collected from participants enrolled to the Pretoria COVID-19 study cohort. Participants were admitted to Tshwane District Hospital (Pretoria, South Africa) between 25 November 2021-20 December 2021 (Table S1). In all waves, samples were collected when more than 90% of SARS-CoV-2 cases in South Africa were caused by the respective variants. Sequence confirmation was only available for a subset of samples but all the samples that were sequenced corresponded to the appropriate variant for that wave. All samples were from HIV-negative individuals who were above 18 years of age and provided consent. Ethical clearance was obtained for each cohort from Human Research Ethics Committees from the University of Pretoria (247/2020) and University of Cape Town (R021/2020). All participants had PCR-confirmed SARS-CoV-2 infection before blood collection Written informed consent was obtained from all participants. BTI participants were recruited from HCWs at the NICD, Steve Biko Academic Hospital (Tshwane, South Africa) and Groote Schuur Hospital (Cape Town, South Africa). Lack of prior infection in these individuals was confirmed by Nucleocapsid ELISA.

#### Cell lines

Human embryo kidney HEK293T cells were cultured at 37°C, 5% CO_2_, in DMEM containing 10% heat-inactivated fetal bovine serum (Gibco BRL Life Technologies) and supplemented with 50 μg/ml gentamicin (Sigma). Cells were disrupted at confluence with 0.25% trypsin in 1 mM EDTA (Sigma) every 48–72 hours. HEK293T/ACE2.MF cells were maintained in the same way as HEK293T cells but were supplemented with 3 μg/ml puromycin for selection of stably transduced cells. HEK293F suspension cells were cultured in 293 Freestyle media (Gibco BRL Life Technologies) and cultured in a shaking incubator at 37°C, 5% CO_2_, 70% humidity at 125rpm maintained between 0.2 and 0.5 million cells/ml. Jurkat-Lucia™ NFAT-CD16 cells were maintained in IMDM media with 10% heat-inactivated fetal bovine serum (Gibco, Gaithersburg, MD), 1% Penicillin Streptomycin (Gibco, Gaithersburg, MD) and 10 μg/ml of Blasticidin and 100 μg/ml of Zeocin was added to the growth medium every other passage. Cells were cultured at 37°C, 5% CO_2_ in RPMI containing 10% heat-inactivated fetal bovine serum (Gibco, Gaithersburg, MD) with 1% Penicillin Streptomycin (Gibco, Gaithersburg, MD) and 2-mercaptoethanol to a final concentration of 0.05 mM and not allowed to exceed 4 x 10^5^ cells/ml to prevent differentiation.

### METHOD DETAILS

#### Spike plasmid and Lentiviral Pseudovirus Production

The SARS-CoV-2 Wuhan-1 spike, cloned into pCDNA3.1 was mutated using the QuikChange Lightning Site-Directed Mutagenesis kit (Agilent Technologies) and NEBuilder HiFi DNA Assembly Master Mix (NEB) to include D614G (original) or lineage defining mutations for Beta (L18F, D80A, D215G, 242-244del, K417N, E484K, N501Y, D614G and A701V), Delta (T19R, 156-157del, R158G, L452R, T478K, D614G, P681R and D950N), Omicron BA.1 (A67V, Δ69-70, T95I, G142D, Δ143-145, Δ211, L212I, 214EPE, G339D, S371L, S373P, S375F, K417N, N440K, G446S, S477N, T478K, E484A, Q493R, G496S, Q498R, N501Y, Y505H, T547K, D614G, H655Y, N679K, P681H, N764K, D796Y, N856K, Q954H, N969K, L981F), Omicron BA.2 (T19I, L24S, 25-27del, G142D, V213G, G339D, S371F, S373P, S375F, T376A, D405N, R408S,K417N,N440K, S477N, T478K, E484A, Q493R, Q498R, N501Y, Y505H, D614G, H655Y, N679K, P681H, N764K, D796Y, Q954H, N969K) or Omicron BA.4 T19I, L24S, Δ25-27, Δ69-70, G142D, V213G, G339D, S371F, S373P, S375F, T376A, D405N, R408S, K417N, N440K, L452R, S477N, T478K, E484A, F486V, Q498R, N501Y, Y505H, D614G, H655Y, N679K, P681H, N764K, D796Y, Q954H, N969K).

Pseudotyped lentiviruses were prepared by co-transfecting HEK293T cell line with the SARS-CoV-2 ancestral variant spike (D614G), Beta, Delta, C.1.2, Omicron BA.1 or Omicron BA.2 spike plasmids in conjunction with a firefly luciferase encoding lentivirus backbone (HIV-1 pNL4.luc) plasmid as previously described^7^. Culture supernatants were clarified of cells by a 0.45-μM filter and stored at −70 °C. Other pcDNA plasmids were used for the ADCC assay.

#### Pseudovirus neutralization assay

For the neutralization assay, plasma samples were heat-inactivated and clarified by centrifugation. Heat-inactivated plasma samples were incubated with the SARS-CoV-2 pseudotyped virus for 1 hour at 37°C, 5% CO_2_. Subsequently, 1×10^4^ HEK293T cells engineered to over-express ACE-2 (293T/ACE2.MF)(kindly provided by M. Farzan (Scripps Research)) were added and incubated at 37°C, 5% CO_2_ for 72 hours upon which the luminescence of the luciferase gene was measured. Titers were calculated as the reciprocal plasma dilution (ID_50_) causing 50% reduction of relative light units. CB6 and CA1 was used as positive controls for D614G, Beta and Delta. 084-7D, a mAb targeting K417N was used as a positive control for Omicron BA.1 and Beta.

#### Antibody-dependent cellular cytotoxicity (ADCC) assay

The ability of plasma antibodies to cross-link and signal through FcγRIIIa (CD16) and spike expressing cells was measured as a proxy for ADCC. For spike assays, HEK293T cells were transfected with 5μg of SARS-CoV-2 spike plasmids using PEI-MAX 40,000 (Polysciences) and incubated for 2 days at 37°C. Expression of spike was confirmed by differential binding of CR3022 and P2B-2F6 and their detection by anti-IgG APC staining measured by flow cytometry. Subsequently, 1×10^5^ spike transfected cells per well were incubated with heat inactivated plasma (1:100 final dilution) or monoclonal antibodies (final concentration of 100 μg/ml) in RPMI 1640 media supplemented with 10% FBS 1% Pen/Strep (Gibco, Gaithersburg, MD) for 1 hour at 37°C. Jurkat-Lucia™ NFAT-CD16 cells (Invivogen) (2×10^5^ cells/well and 1×10^5^ cells/well for spike and other protein respectively) were added and incubated for 24 hours at 37°C, 5% CO_2_. Twenty μl of supernatant was then transferred to a white 96-well plate with 50 μl of reconstituted QUANTI-Luc secreted luciferase and read immediately on a Victor 3 luminometer with 1s integration time. Relative light units (RLU) of a no antibody control was subtracted as background. Palivizumab was used as a negative control, while CR3022 was used as a positive control, and P2B-2F6 to differentiate the Beta from the D614G variant. 084-7D was used as a positive control for Omicron BA.1 and Beta. To induce the transgene 1x cell stimulation cocktail (Thermofisher Scientific, Oslo, Norway) and 2 μg/ml ionomycin in R10 was added as a positive control to confirm sufficient expression of the Fc receptor. RLUs for spikes were normalised to each other and between runs using CR3022. All samples were run head to head in the same experiment as were all variants tested.

### QUANTIFICATION AND STATISTICAL ANALYSIS

Analyses were performed in Prism (v9; GraphPad Software Inc, San Diego, CA, USA). Non-parametric tests were used for all comparisons. The Mann-Whitney and Wilcoxon tests were used for unmatched and paired samples, respectively. The Friedman test with Dunns correction for multiple comparisons was used for matched comparisons across variants. All correlations reported are non-parametric Spearman’s correlations. *P* values less than 0.05 were considered to be statistically significant.

